## Supplementary Figures for "Phosphoglycerate mutase regulates Treg differentiation through control of serine synthesis and one-carbon metabolism"

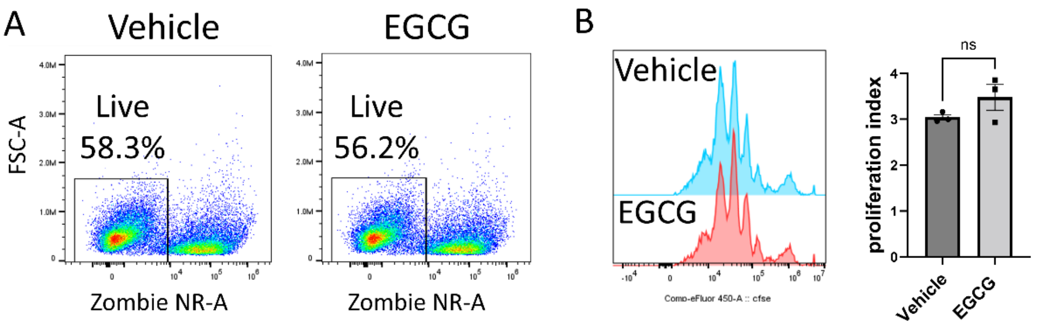


**Figure S1. EGCG has no effect on Treg viability or proliferation**. Naïve murine CD4 cells were cultured under Treg polarizing conditions with either vehicle or EGCG (20 μM). (**A**) Viability was assayed with zombie NIR dye. (**B**) Cell proliferation was assayed via cell proliferation dye, with proliferation index calculated as the total number of divisions divided by the number of cells that went into division. Data derived from 3 biological replicates. ns=not-significant by student's t-test.

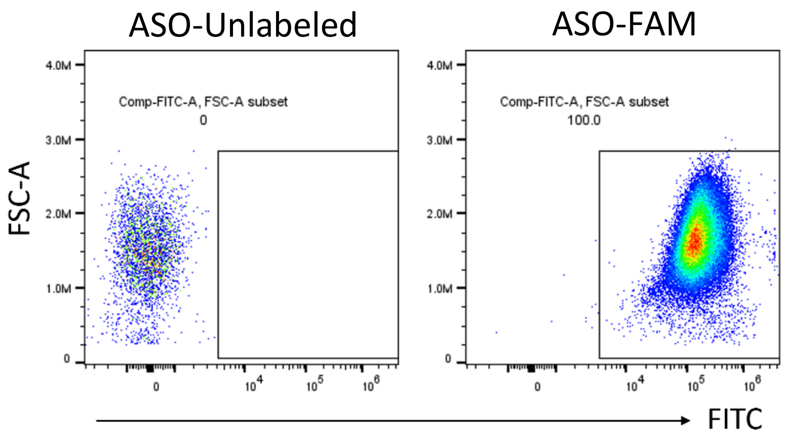


**Figure S2. Efficient uptake of ASOs by cultured CD4 cells**. Naïve murine CD4 cells were cultured under Treg polarizing conditions for 24 hours with scrambled ASOs that were either unlabeled or labeled with a fluorescein (FAM) tag (detected in the FITC channel). Uptake of ASOs was assessed by measuring FITC signal within cells via flow cytometry.

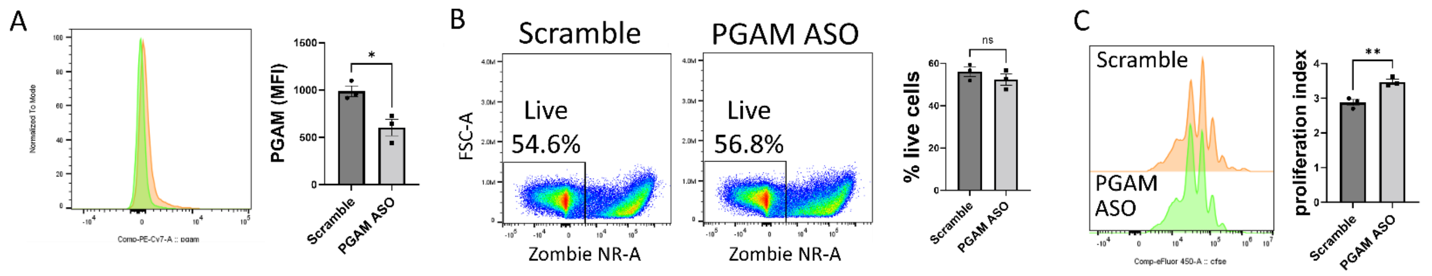


**Figure S3. PGAM ASOs reduce PGAM expression without impacting viability**. Naïve murine CD4 cells were cultured under Treg polarizing conditions for 72 hours with either scrambled or anti-PGAM ASOs. (**A**) PGAM1 expression and (**B**) cell viability were assayed by flow cytometry. Data represent mean ± SEM from 3 biological replicates. (**C**) Cell proliferation dye was used to quantify proliferation index, calculated as in figure S1B. Data represent mean ± SEM from 3 biological replicates. **P* < 0.05, ** P< 0.01 or ns=not-significant by student's t-test.

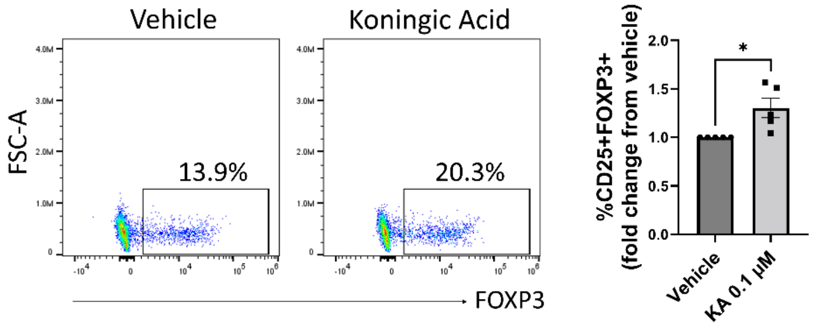


**Figure S4**. **GAPDH inhibition promotes Treg polarization**. Naïve murine CD4 cells were cultured under Treg polarizing conditions with vehicle or koningic acid (KA, 0.1 μM) for 96 hours. Treg polarization was assayed by flow cytometry. Data represent mean ± SEM from 5 independent experiments performed in triplicate. * P <0.05 by Mann-Whitney U test.

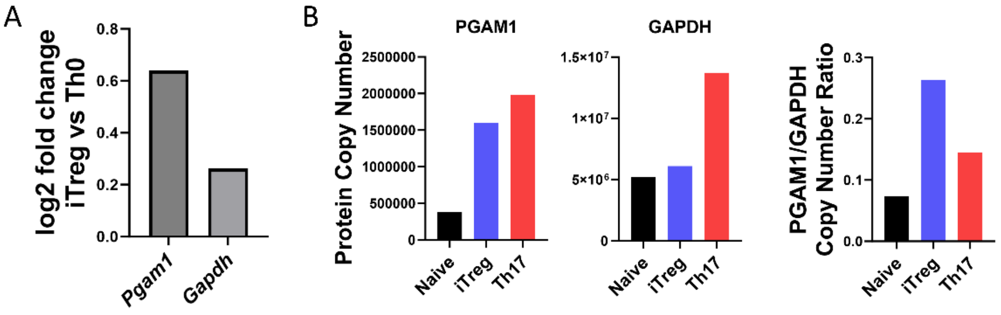


**Figure S5. Increased PGAM to GAPDH expression ratio in regulatory T cells**. (**A**) Analysis of a publicly available RNAseq dataset from human naïve CD4 cells cultured under either Th0 or Treg polarizing conditions for 72 hours (Ubaid Ullah et al., 2018) shows log2 fold change of *Pgam1* expression versus *Gapdh* expression in iTreg cells. (**B**) Data from the ImmPres immunological proteome resource (Brenes et al., 2023) showing protein copy number of PGAM1 (*left*) and GAPDH (*middle*) as well as the ratio of PGAM1 to GAPDH protein copy number (*right*), derived from murine CD4 cells in the naïve state or polarized under Treg or Th17 conditions.

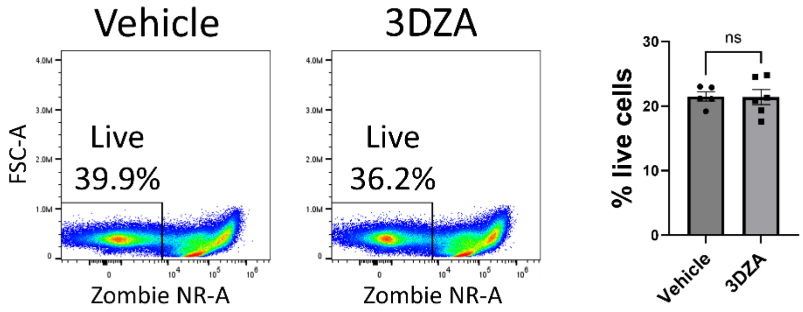


**Figure S6. 3DZA does not affect viability**. Naïve murine CD4 cells were cultured under Treg polarizing conditions for 72 hours with either vehicle or 3DZA (5 μM). Cell viability was assayed by flow cytometry. Data represent mean ± SEM from 5-6 biological replicates. ns=not-significant by student's t-test.

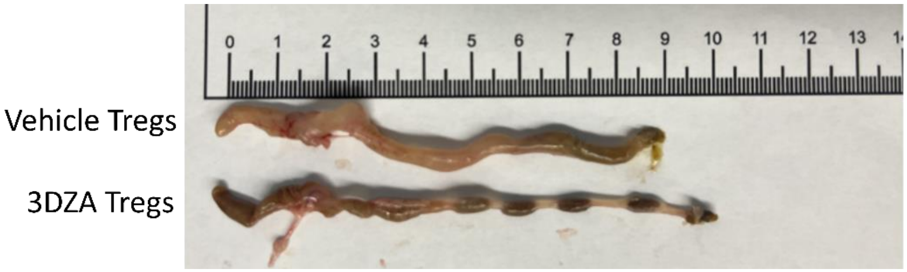


**Figure S7. Tregs polarized with 3DZA are more suppressive in the T cell transfer model of autoimmune colitis**. Naïve murine CD4 T cells were cultured under Treg polarizing conditions with either vehicle or 3DZA and $10^6$ cells were transferred into $RAG^{-/-}$ mouse recipients along with $10^7$ activated CD4 T cells. Mice injected with vehicle-treated Tregs showed colonic wall thickening indicative of more severe colitis.
